# SETDB1 regulates microtubule dynamics

**DOI:** 10.1101/2021.05.15.444210

**Authors:** Rosari Hernandez-Vicens, Nomi Pernicone, Tamar Listovsky, Gabi Gerlitz

**Affiliations:** Department of Molecular Biology, Faculty of Life Sciences and Ariel Center for Applied Cancer Research, Ariel University, Ariel 40700, Israel; Adelson School of Medicine, Ariel University, Ariel 40700, Israel

**Keywords:** Microtubules, tubulin acetylation, lysine methyltransferases, mitosis, HDAC6

## Abstract

SETDB1 is a methyltransferase responsible for the methylation of histone H3-lysine-9, which is mainly related to heterochromatin formation. SETDB1 is overexpressed in various cancer types and is associated with an aggressive phenotype. In agreement with its activity, it mainly exhibits a nuclear localization; however, in several cell types a cytoplasmic localization was reported. Here we show that a substantial cytoplasmic pool of SETDB1 is colocalized with microtubules. Significantly, silencing of SETDB1 led to faster polymerization and reduced rate of catastrophe events of microtubules in parallel to reduced proliferation rate and slower mitotic kinetics. Interestingly, over-expression of either wild-type or catalytic dead SETDB1 altered microtubule polymerization rate to the same extent, suggesting that SETDB1 affects MT dynamics by a methylation-independent mechanism. Finding interaction between SETDB1 and the tubulin deacetylase HDAC6 and increased tubulin acetylation levels upon silencing of SETDB1 suggest a model in which SETDB1 affects microtubule dynamics by interacting with both microtubules and HDAC6 to enhance tubulin deacetylation. Overall, our results suggest a novel cytoplasmic role for SETDB1 in the regulation of microtubule dynamics.

## Introduction

SETDB1 (also known as ESET and KMT1E) is a lysine methyltransferase (KMT) that belongs to the SUV39 family of KMTs that mainly methylate lysine 9 in histone H3 (H3K9). The SUV39 family members are characterized by cysteine-rich pre- and post-SET domains flanking a central SET domain that is responsible for the catalytic activity (Mozzetta et al., 2015; Torrano et al., 2019). SETDB1 also contains a methyl-CpG-binding domain (MBD) and a triple Tudor domain responsible for binding of H3K9me/K14ac to promote histone deacetylation by HDACs (Jurkowska et al., 2017; Schultz, 2002). SETDB1 can mono-, di- and tri-methylate H3K9. H3K9me2/3 are usually associated with gene repression and heterochromatin formation (Mozzetta et al., 2015), and indeed SETDB1 is involved in silencing the X chromosome (Keniry et al., 2016; Minkovsky et al., 2014), repetitive elements (Cuellar et al., 2017; Liu et al., 2014; Matsui et al., 2010; Sharif et al., 2016) and specific genes (Du et al., 2018; Jiang et al., 2010; Karimi et al., 2011). SETDB1 is important for various developmental processes including embryogenesis (Bilodeau et al., 2009; Dodge et al., 2004; Zeng et al., 2019), neurogenesis (Jiang et al., 2017), immune cell development (Hachiya et al., 2016; Takikita et al., 2016), germ line development (Liu et al., 2014) and chondrocyte differentiation (Lawson et al., 2013). SETDB1 was also shown to methylate non-histone proteins such as ING2 (Binda et al., 2010), p53 (Fei et al., 2015), UBF (Hwang et al., 2014) and Akt (Guo et al., 2019; Wang et al., 2019).

SETDB1 methylation of H3K9, as well as most of its non-histone proteins that are nuclear proteins, is in correlation with its nuclear localization. However, a cytoplasmic pool of SETDB1 has been found in several types of cells including HeLa cells (Tachibana et al., 2015), HEK293 cells (Wang et al., 2019), mouse embryonic fibroblasts (Cho et al., 2013), differentiated myoblasts (Beyer et al., 2016) and human melanoma biopsies (Kostaki et al., 2014). Cytoplasmic localization of SETDB1 is thought to facilitate methylation of newly synthesized histones before their incorporation into nucleosomes (Loyola et al., 2006) or to restrict the enzyme of methylating nucleosomal H3K9 (Beyer et al., 2016; Cho et al., 2013).

In recent years SETDB1 has been considered as an oncogene. Its genomic location is commonly amplified in melanoma (Ceol et al., 2011; Orouji et al., 2019) and its expression levels are increased in various types of cancer including melanoma (Ceol et al., 2011; Orouji et al., 2019), colorectal cancer (Chen et al., 2017; Ho et al., 2017; Yu et al., 2019), liver cancer (Fei et al., 2015; Wong et al., 2016; Zhang et al., 2018) and lung cancer (Cruz-Tapias et al., 2019; Kang and Min, 2020; Rodriguez-Paredes et al., 2014; Sun et al., 2015). SETDB1 oncogenic behavior supports cancer cell proliferation, migration and invasion (Chen et al., 2017; Fei et al., 2015; Gauchier et al., 2019; Guo et al., 2019; Orouji et al., 2019; Rodriguez-Paredes et al., 2014; Sun et al., 2015; Wang et al., 2019; Wong et al., 2016; Yu et al., 2019; Zhang et al., 2018). More recently SETDB1 was also linked to adaptive resistance of tumor cells to various drugs (Rodriguez-Paredes et al., 2014; Torrano et al., 2019) and repression of the innate immune response (Cuellar et al., 2017; Kang, 2018).

Currently, SETDB1 is thought to promote cancer mainly by its nuclear activity of methylating H3K9 or transcription factors such as p53 (Fei et al., 2015). Since a substantial amount of SETDB1 can be found in the cytoplasm, we wondered whether SETDB1 has a specific cytoplasmic activity that may also promote cancer formation and progression. Here we show that cytoplasmic SETDB1 co-localizes with the MT network. SETDB1 knockdown (KD) increased MT stability as measured by EB1-tracking and MT recovery from nocodazole treatment and interfered with mitotic progression. Over-expression of either wild type or catalytic dead (CD) SETDB1 increased MT polymerization rate to the same extent, suggesting SETDB1 affects MT dynamics in a methyltransferase-independent manner. Finding interaction between SETDB1 and the tubulin deacetylase HDAC6, along with increased tubulin acetylation levels after knockdown of SETDB1, suggest that SETDB1 can affect MT dynamics by supporting HDAC6 activity.

## Results

### SETDB1 associates with the MT network

To identify a cytoplasmic role for SETDB1, we first verified its cytoplasmic localization in both mouse and human melanoma cells: B16-F1 and WM266.4 cells, respectively. Biochemical fractionation followed by Western blot analysis identified a substantial amount of SETDB1 in the cytoplasmic fraction of these cells in contrast to another H3K9 methyltransferase, SUV39H2, which was found only in the nuclear fraction (Fig 1A). Immunostaining for SETDB1 verified this observation and revealed partial co-localization of SETDB1 with MTs (Fig 1B,C). To validate the immunostaining we used a different antibody against SETDB1 that revealed a similar pattern of localization in human WM266.4 cells (Fig 1D,E). Co-localization of SETDB1 with microtubules was also found in HeLa cells (Fig 1F). Moreover, this pattern was kept during mitosis as well (Fig 1G). To verify these results, we analyzed cells over-expressing GFP-fused SETDB1 and revealed a similar pattern of partial co-localization with MTs (Fig1 H,I). Hence our results present SETDB1 association with MTs in several cell types both in interphase and mitosis suggesting a role for SETDB1 in MT functions.

**Figure 1.**
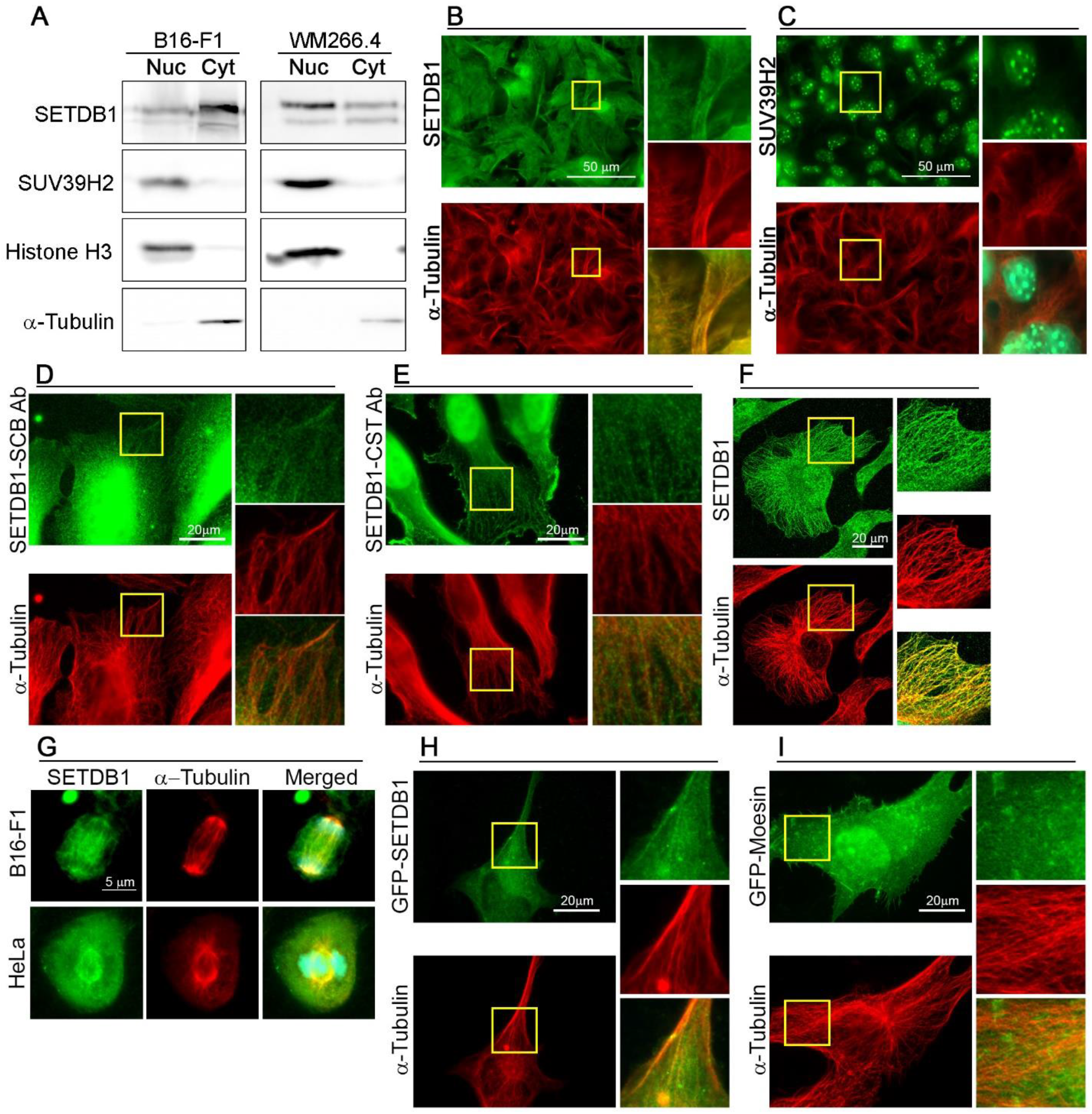
SETDB1 is found in the cytoplasm and partially co-localizes with MTs. **A** Western blot analysis of nuclear (Nuc) and cytoplasmic (Cyt) fractions of mouse B16-F1 cells and human WM266.4 cells for SETDB1, SUV39H2, α-Tubulin and histone H3. **B**,**C** B16-F1 cells immunostained for SETDB1 or SUV39H2 and α-Tubulin. DNA stained with Hoechst 33342. Sections inside the inserts are magnified at the right side of each micrograph. The merged images show the merged signals of SETDB1, α-Tubulin and Hoechst. Scale bar: 50 μm. **D**,**E** WM266.4 cells immunostained for SETDB1 with two different commercial antibodies (SCB=Santa Cruz Biotechnology sc-66884, CST=Cell Signaling Technology 93212) and α-Tubulin. Sections inside the inserts are magnified at the right side of each micrograph. The merged images show the merged signals of SETDB1 and α-Tubulin. Scale bar: 20 μm. **F** HeLa cells immunostained for SETDB1 and α-Tubulin. The merged images show the merged signals of SETDB1 and α-Tubulin. Scale bar: 20 μm. **G** Mitotic B16-F1 cells and HeLa cells immunostained for SETDB1 and α-Tubulin. DNA stained with Hoechst 33342. Sections inside the inserts are magnified at the right side of each micrograph. The merged images show the merged signals of SETDB1, α-Tubulin and Hoechst. Scale bar: 5 μm. **H**,**I** WM266.4 cells transfected with either pEGFP-Moesin (H) or pEGFP-SETDB1 (I) immunostained for GFP and α-Tubulin. Sections inside the inserts are magnified at the right side of each micrograph. The merged images show the merged signals of GFP and α-Tubulin. Scale bar: 20 μm.

### SETDB1 affects MT growth rate

To evaluate whether SETDB1 can affect MT functions, we first tested the impact of SETDB1 silencing on the rate of MT growth during recovery from nocodazole treatment. B16-F1 cells were transfected with control siRNA or SETDB1 siRNA (Sup Fig 1) and treated with nocodazole for 3 hours to depolymerize MTs. Following nocodazole washout, the cells were further incubated for an additional three or seven minutes, fixed and immunostained for α-tubulin. Measurements of the area covered by MTs originated from the MTOC revealed an increase of 82 % and 60 % in SETDB1 KD cells, in comparison to control cells at the three minutes and seven minutes time points, respectively (Fig 2). This finding suggests that SETDB1 attenuates MT growth rate.

**Figure 2.**
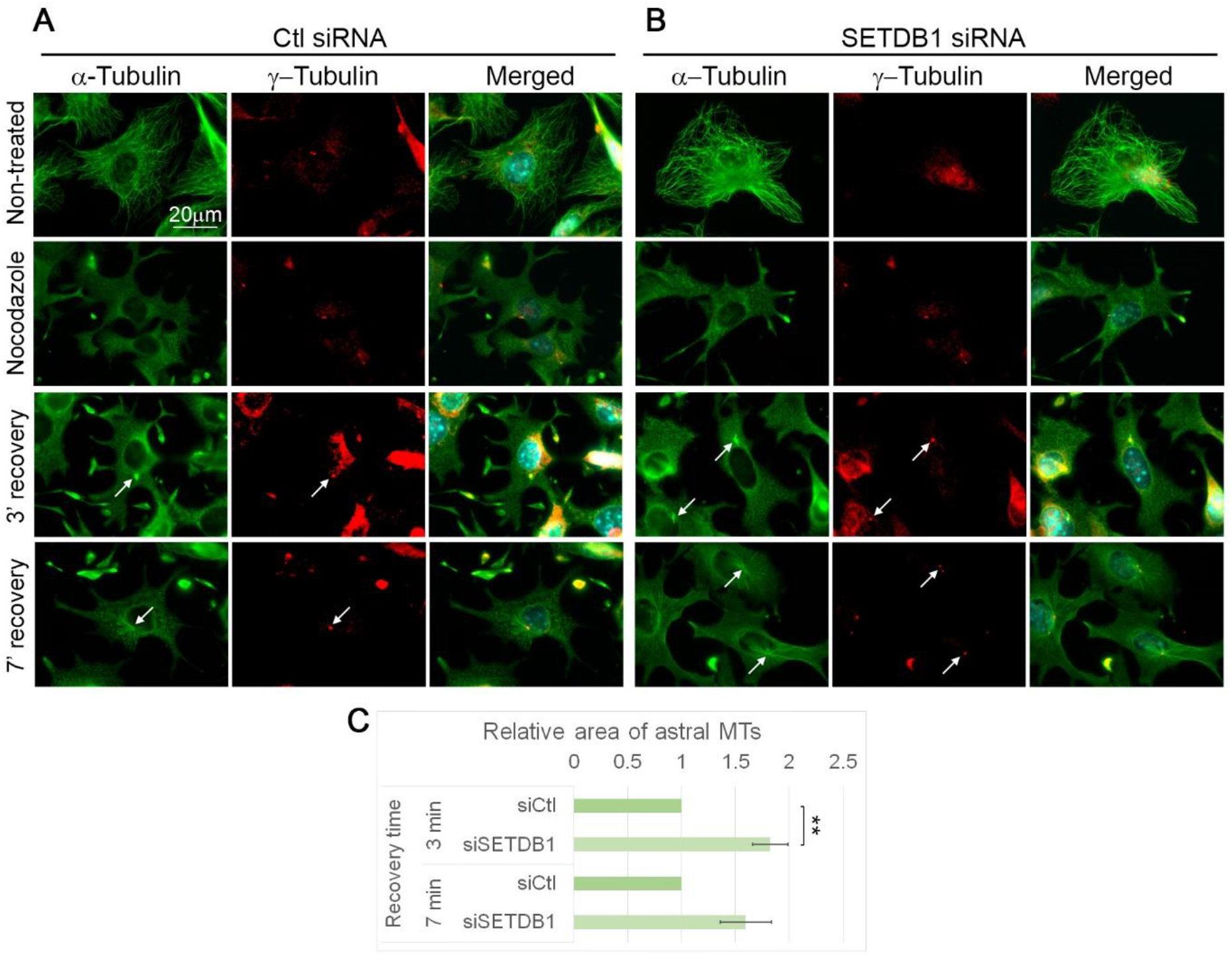
SETDB1 KD enhances MT polymerization rate. **A**,**B** MTs were disrupted in B16-F1 cells transfected with either Ctl siRNA (A) or siRNA against SETDB1 (B) by nocodazole treatment for 3 hours. Following nocodazole removal, cells were further incubated for the indicated time periods to allow MT polymerization. After fixation, cells were immunostained with antibodies against α-Tubulin, γ-Tubulin and DNA was stained with Hoechst 33342. The white arrows indicate the localization of the MTOCs. Scale bar: 20 μm. **C** Quantification of the recovery rate of MTs after nocodazole washout. The area covered by MTs around the MTOC was quantified by ImageJ software and normalized to the area in cells transfected with ctl siRNA. The bar graph shows the average of the relative area covered by MTs of three repetitions ± SE. At least of 30 cells were measured for each condition in each repetition. Statistical significance was calculated with Student’s *t-*test, ** *P* < 0.01.

SETDB1 KD reduces the protein levels in both the cytoplasm and the nucleus; therefore, the KD effects may be due to reduced nuclear activity of SETDB1 as a transcriptional regulator or due to reduced association of cytoplasmic SETDB1 with the MT network. Notably, over-expressed (OE) SETDB1 is localized to the cytoplasm both in previous reports (Cho et al., 2013; Tachibana et al., 2015; Tsusaka et al., 2019) and in our hands (Fig 1H, 3B). Therefore, we repeated the MT recovery assay after nocodazole treatment with cells over-expressing GFP-SETDB1 or GFP-Moesin that served as a negative control. As shown in Fig 3, the levels of MTOC-linked MTs 3 minutes after nocodazole washout in GFP-SETDB1 OE cells were reduced by half, compared to GFP-Moesin OE cells. This effect was in the opposite direction to the SETDB1 silencing effect (Fig 2), suggesting that the SETDB1 effect on MT growth is due to the activity of its cytoplasmic pool rather than the nuclear SETDB1.

**Figure 3.**
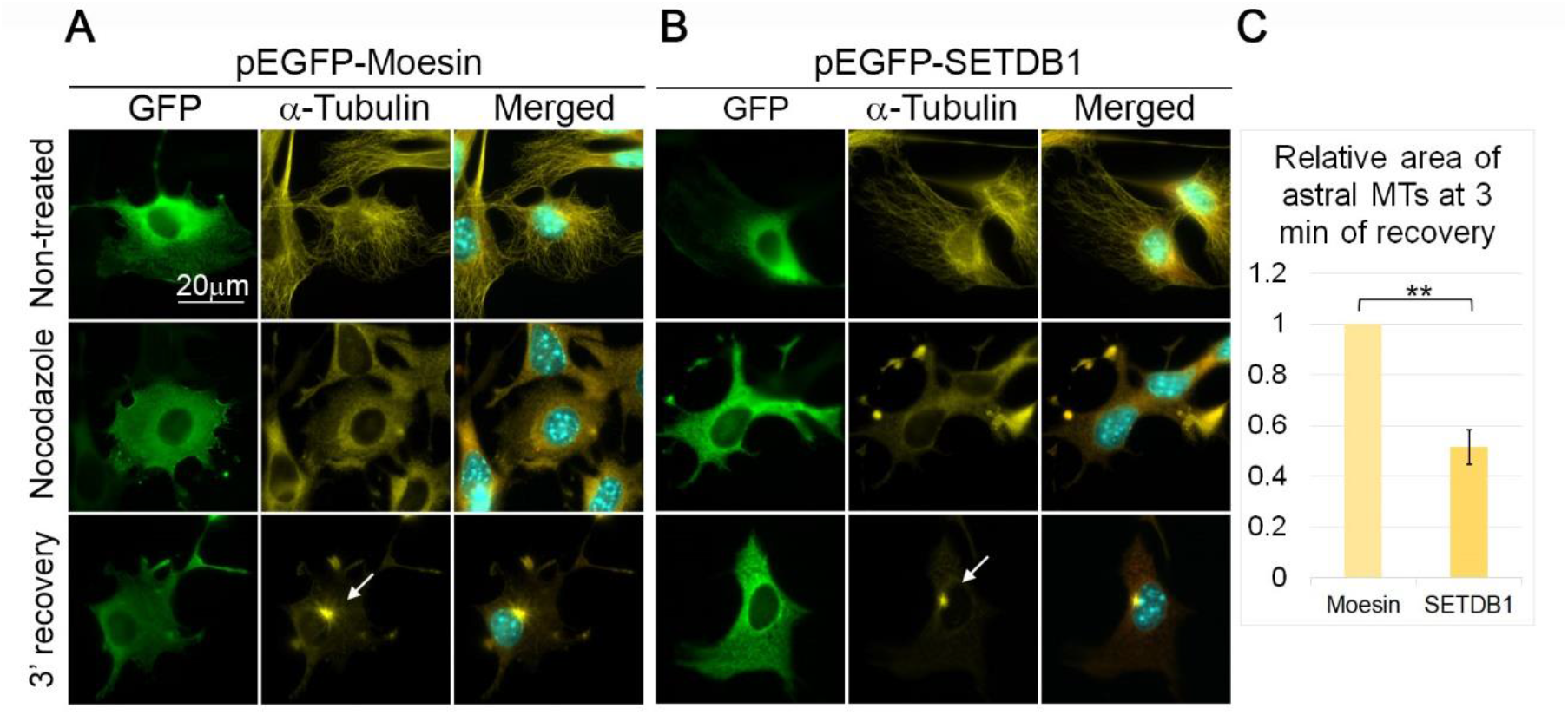
SETDB1 over-expression enhances MT polymerization rate. **A**,**B** MTs were disrupted in B16-F1 cells transfected with either pEGFP-Moesin (A) or pEGFP-SETDB1 (B) by nocodazole treatment for 3 hours. Following nocodazole removal, cells were further incubated for 3 minutes to allow MT polymerization. After fixation, cells were immunostained with antibodies against GFP and α-Tubulin and DNA was stained with Hoechst 33342. The white arrows indicate the localization of the MTOCs. Scale bar: 20 μm. **C** Quantification of the recovery rate of MTs after nocodazole washout. The area covered by MTs around the MTOC was quantified by ImageJ software and normalized to the area in cells transfected with pEGFP-Moesin. The bar graph shows the average of the relative area covered by MTs of three repetitions ± SE. At least of 30 cells were measured for each condition in each repetition. Statistical significance was calculated with Student’s *t-*test, ** *P* < 0.01.

To verify the effects of SETDB1 on MT growth we monitored the MT plus end dynamics by tracking GFP-fused EB1 in SETDB1 KD cells (Sup Fig 1, Fig 4). Notably, MT growth duration, growth length and growth rate were increased in SETDB1 KD cells by 25%, 38% and 11%, respectively, in comparison to control cells (Fig 4B-D). On the other hand, MT catastrophe rate in SETDB1 KD cells was reduced by 15% in comparison to control cells (Fig 4E). These results suggest that SETDB1 is a negative regulator of MT growth.

**Figure 4.**
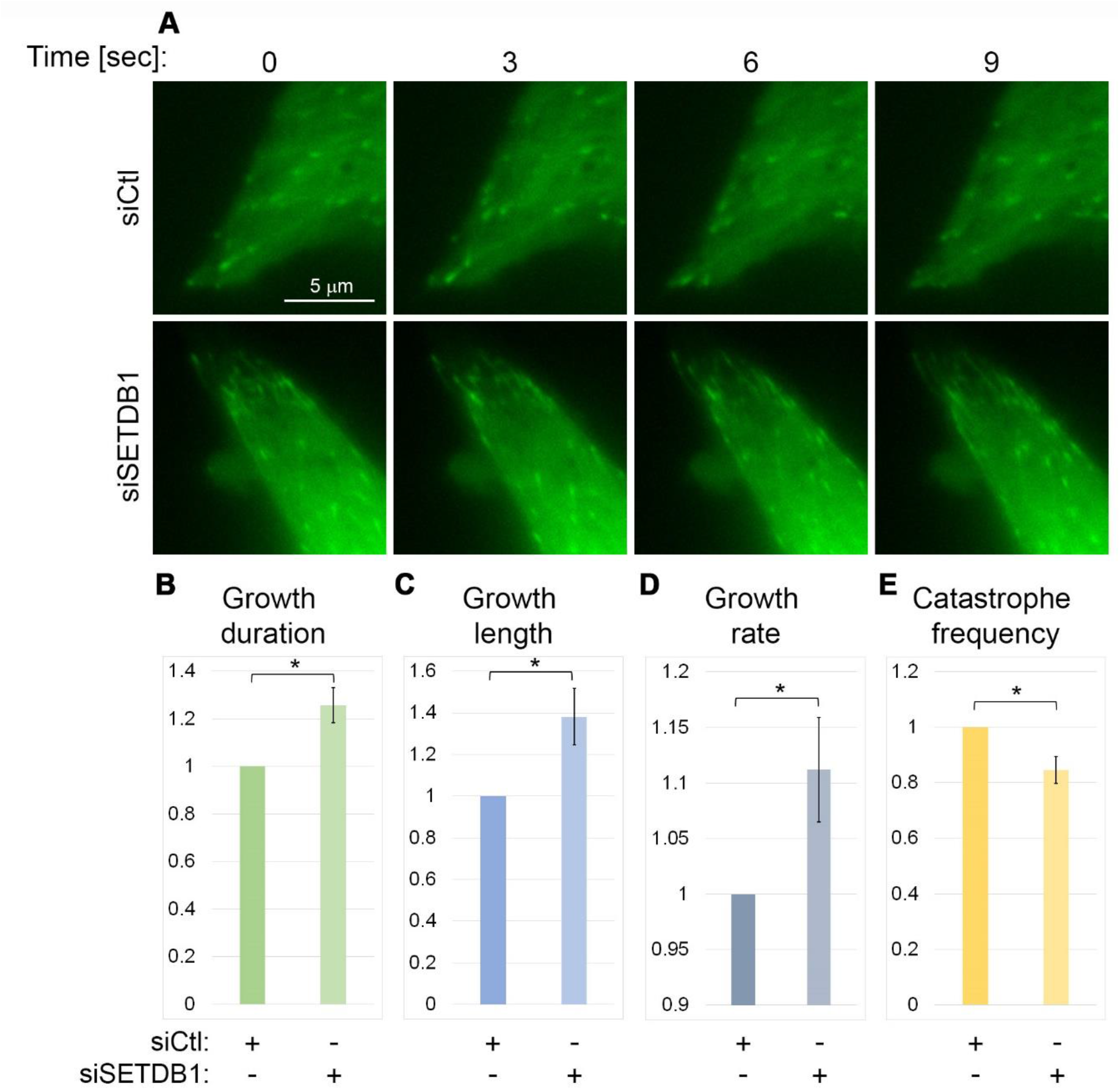
SETDB1 KD affects MT dynamics. **A** Micrographs showing signals of GFP comets in HeLa cells co-transfected with EB1-GFP vector and either Ctl siRNA or siRNA against SETDB1. Frames were taken every 3 seconds. Scale bar: 5 μm. **B-E** Parameters of MT dynamics as measured by EB1 tracking in time lapse microscopy. The bar charts represent the ratios of the indicated parameters of MTs in SETDB1 KD cells to Ctl cells. The averages are of three repetitions ± SE. In each experiment a minimum of 100 EB1-GFP comets per condition were measured. Statistical significance was calculated with Student’s *t*-test, * *P* < 0.05.

### SETDB1 is important for proper cell division

The finding that SETDB1 regulates MT dynamics led us to test if SETDB1 is important for cell division, a process that is heavily dependent on MT organization. Indeed, SETDB1 KD attenuated the cellular proliferation rate by 20% (Fig 5A), while increasing unsuccessful mitotic events from 4.1% to 22.1% (Fig 5B). Measuring the lengths of the different cell division stages in cells that were able to finish mitosis successfully revealed a significant increase in the duration of both early and late mitosis in SETDB1 KD cells in comparison to control cells: in SETDB1 KD cells the durations from nuclear envelope breakdown (NEB) to anaphase and from anaphase to the appearance of a cleavage furrow were increased by 17% and 40%, respectively, in comparison to control cells (Fig 5C-H).

**Figure 5.**
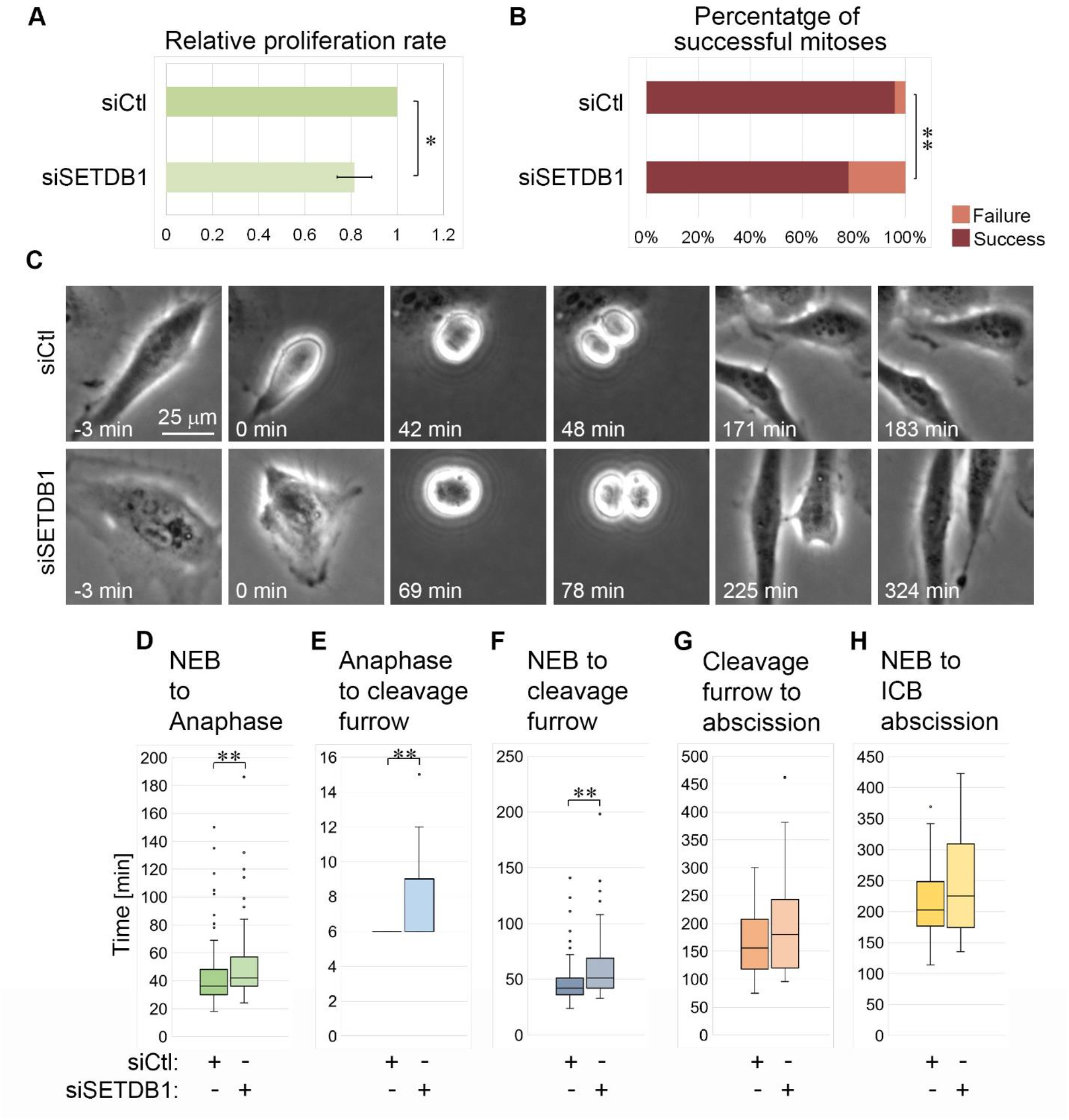
SETDB1 KD alters the duration of mitotic phases. **A** Relative proliferation rate of SETDB1 KD HeLa cells as measured by the XTT assay. The average of three repetitions ± SE is shown. Statistical significance was calculated with Student’s *t*-test, * *P* < 0.05. **B** The percentage of successful and failed mitotic event in HeLa cells transfected with either Ctl siRNA or siRNA against SETDB1. Statistical significance was calculated with Student’s *t*-test, ** *P* < 0.01. **C** Micrographs showing mitotic progression of HeLa cells transfected with either Ctl siRNA or siRNA against SETDB1. Scale bar: 25 μm. **D-H** Time periods of the indicated phases during cell division as calculated from time-lapse images of HeLa cells transfected with either Ctl siRNA or siRNA against SETDB1. Frames were taken every 3 minutes. In each experiment a minimum of 40 cells were tracked for each condition. The average values of a representative experiment out of three experiments are presented. Statistical significance was calculated with Wilcoxon-Mann-Whitney test, ** *P* < 0.01.

### SETDB1 affects MT growth rate independently of its methyltransferase activity

SETDB1 is a well-established methyltransferase that was found to methylate both histones (Schultz, 2002) and non-histone proteins (Fei et al., 2015; Guo et al., 2019; Wang et al., 2019). Moreover, recently α-tubulin was found to be methylated at lysine 40 by SETD2 (Park et al., 2016). To evaluate if the effect of SETDB1 on MT growth rate is dependent on its methyltransferase activity we repeated the MT recovery assay while over-expressing either WT SETDB1 or H1224K point mutated SETDB1 (Fig 6). The H1224K mutation was shown before to completely impair the methyltransferase activity of SETDB1 (Schultz, 2002). As shown in Fig. 6, over-expression of the enzymatically inactive H1224K SETDB1 attenuated MT recovery from nocodazole treatment to the same extent as over-expression of WT SETDB1. Thus, SETDB1 can affect MT dynamics by a mechanism that is independent of its enzymatic activity.

**Figure 6.**
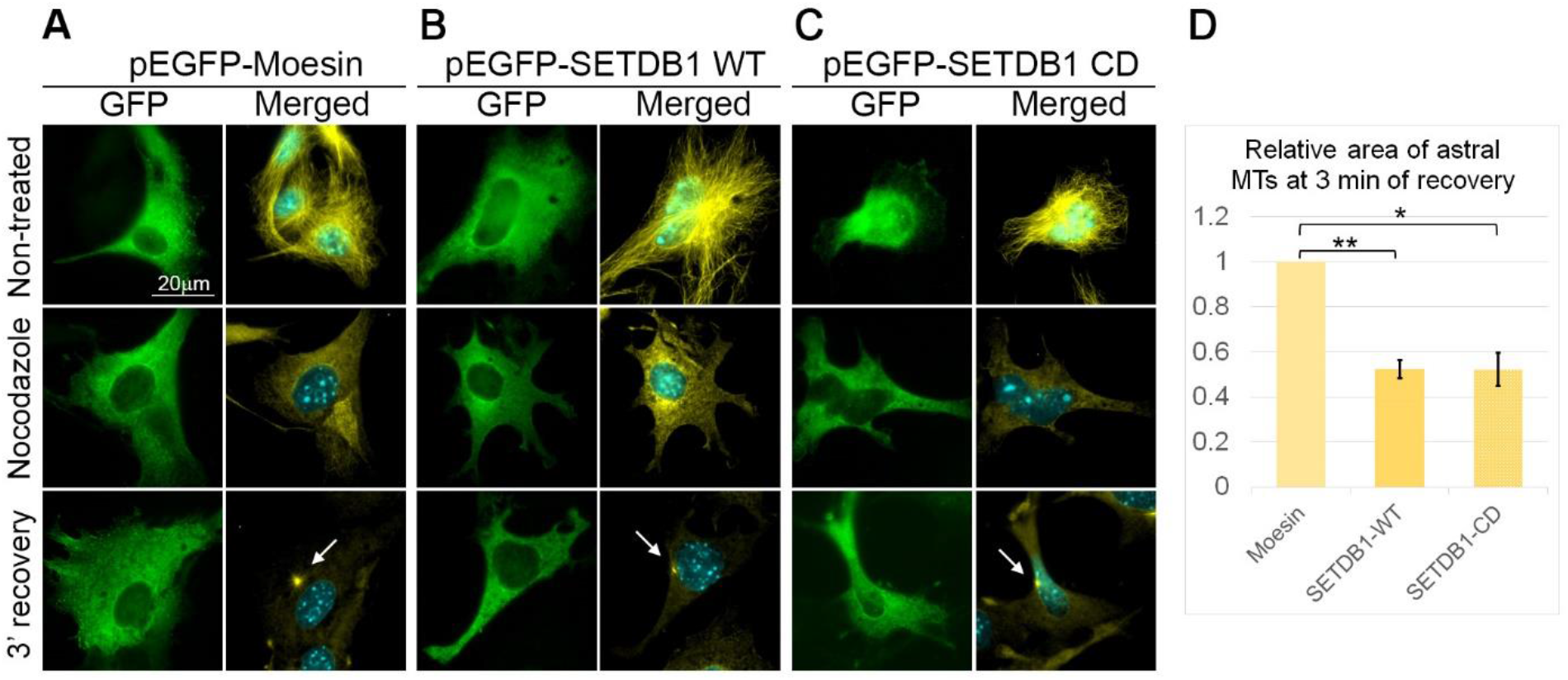
SETDB1 over-expression enhances MT polymerization rate in KMT-independent manner. **A-C** MTs were disrupted in B16-F1 cells transfected with either pEGFP-Moesin (A) or pEGFP-SETDB1 WT (B) or pEGFP-SETDB1 CD (C) by nocodazole treatment for 3 hours. Following nocodazole removal, cells were further incubated for 3 minutes to allow MT polymerization. After fixation, cells were immunostained with antibodies against GFP and α-Tubulin and DNA was stained with Hoechst 33342. The white arrows indicate the localization of the MTOCs. Scale bar: 20 μm. **C** Quantification of the recovery rate of MTs after nocodazole washout. The area covered by MTs around the MTOC was quantified by ImageJ software and normalized to the area in cells transfected with pEGFP-Moesin. The bar graph shows the average of the relative area covered by MTs of three repetitions ± SE. At least of 30 cells were measured for each condition in each repetition. Statistical significance was calculated with Student’s *t-*test, * *P* < 0.05, ** *P* < 0.01.

### SETDB1 affects tubulin acetylation

The molecular mechanism by which SETDB1 affects MT dynamics seemed to be independent of its methyltransferase activity. To identify a molecular mechanism by which SETDB1 can affect MT dynamics we took into account two features: the first is that nuclear SETDB1 is known to participate in a complex with HDAC1 and HDAC2 to repress transcription (Yang et al., 2003). The second is that acetylation of Lys40 in α-tubulin (acetylated tubulin) is a key tubulin post-translational modification, which is associated with stable MTs in various cellular contexts (A et al., 2020; Li and Yang, 2015). We hypothesized that SETDB1 may interact with a tubulin HDAC and affect its activity. Since the major tubulin deacetylase is HDAC6 we tested if SETDB1 can interact with it and affect tubulin acetylation levels. As shown in Fig 7A, HDAC6 co-immunoprecipitated with SETDB1, suggesting an *in vivo* interaction between these two factors. In addition, SETDB1 KD led to an increase of 110% in the levels of tubulin acetylation (Fig 7B,C), thus supporting the hypothesis that SETDB1 can serve as a co-factor of HDAC6 in the cytoplasm to support tubulin deacetylation to regulate MT dynamics.

**Figure 7.**
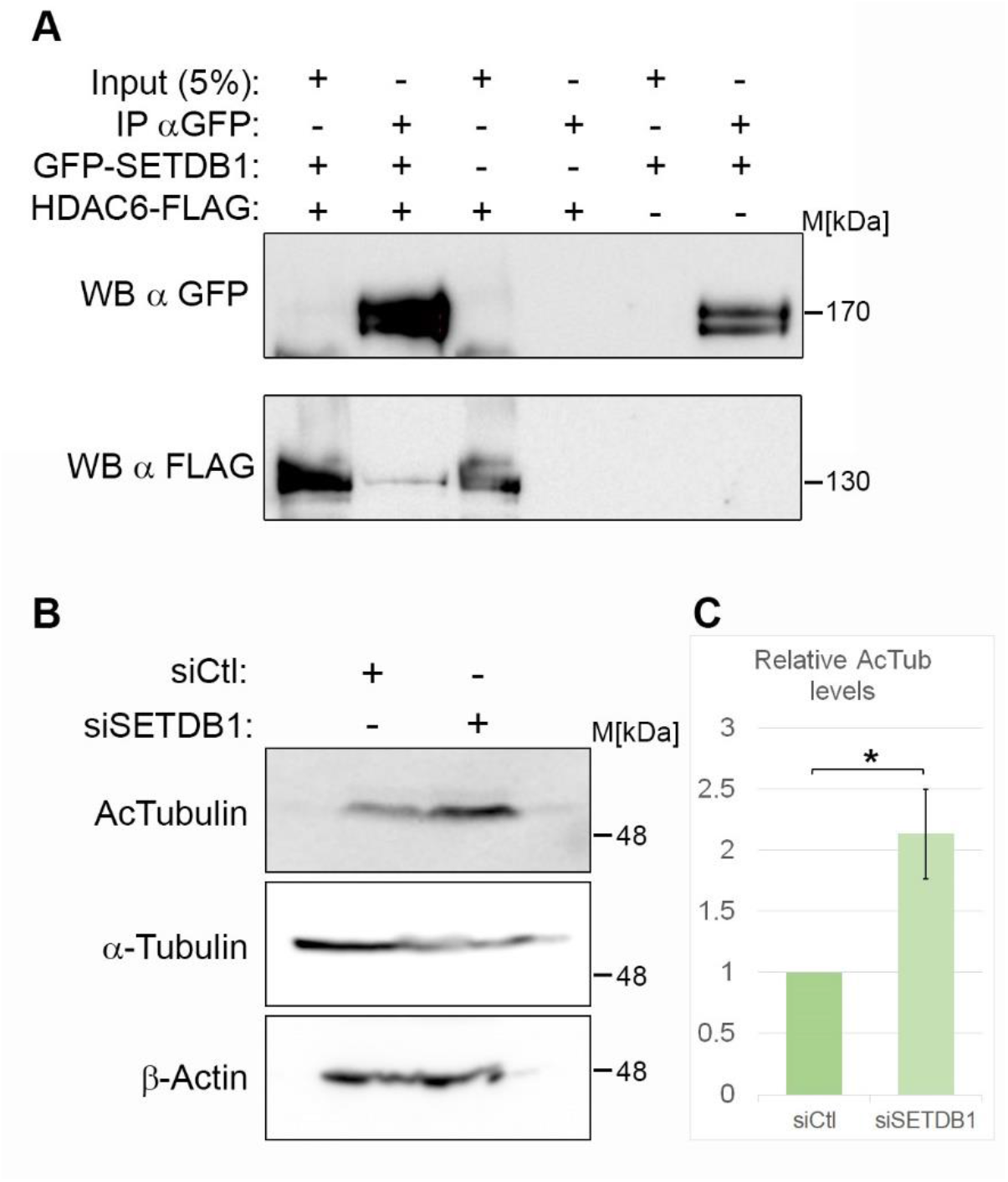
SETDB1-HDAC6 interaction. **A** Co-IP of over-expressed SETDB1-GFP and HDAC6-FLAG in HEK293 cells. **B-C** Western blot analysis of the Ac-Tubulin levels in B16-F1 cells tranfected with either Ctl siRNA or siRNA against SETDB1 (B). The ratio of Ac-Tubulin to Tubulin levels in three repetitions normalized to the same ratio in Ctl siRNA-transfected cells +/-SE. Statistical significance was calculated with Student’s *t-*test, * *P* < 0.05 (C).

## Discussion

SETDB1 is a well-established H3K9 methyltransferase involved in embryogenesis and development (Bilodeau et al., 2009; Dodge et al., 2004; Hachiya et al., 2016; Jiang et al., 2017; Lawson et al., 2013; Liu et al., 2014; Takikita et al., 2016; Zeng et al., 2019) as well as in the etiology of cancer (Batham et al., 2019; Kang, 2018; Karanth et al., 2017; Strepkos et al., 2020). Current perception is that SETDB1 affects these processes due to its nuclear localization, while any cytoplasmic localization of SETDB1 serves to either sequester it or methylate newly generated histone H3 (Beyer et al., 2016; Cho et al., 2013; Loyola et al., 2006). Here we found that cytoplasm localized SETDB1 is partially co-localized with MTs in both mouse and human cells. Moreover, SETDB1 KD altered MT dynamics: it sped up MT recovery from nocodazole treatment and in steady-state conditions it led to enhanced MT growth duration, length and rate in parallel to reduced frequency of catastrophe events. Since SETDB1 KD is not cytoplasmic-specific, we manipulated SETDB1 also by over-expressing it. Notably, over-expressed SETDB1 was localized specifically to the cytoplasm, as was previously found by others. This manipulation led to slower MT recovery from nocodazole treatment, suggesting SETDB1 did not affect MTs by altering the cellular transcriptome, but rather by a more direct mechanism.

Recently, the methyltransferases SETD2 and SET8 were shown to methylate α-tubulin on K40 and K311, respectively, while SETD2 activity was limited to the M phase (Chin et al., 2020; Park et al., 2016). To assess if SETDB1 may affect MT dynamics by its methyltransferase activity, we evaluated the effects of CD SETDB1 on MT dynamics and found that both WT and CD SETDB1 reduced MT recovery rate from nocodazole treatment. These results suggest that the effects of SETDB1 on MTs were methyltransferase independent. This observation is in agreement with studies on SETDB1 oncogenic activity in the zebrafish model: over-expression of SETDB1 was found to accelerate melanoma onset in a zebrafish model for melanoma formation and progression. Notably, over-expression of the enzymatically inactive H1224K SETDB1 (CD SETDB1) still significantly accelerated melanoma onset, although to a lower extent than WT SETDB1 (Ceol et al., 2011). This suggests that the oncogenic function of SETDB1 is partially contributed by methyltransferase independent activities of the protein. One of these activities may be modulation of MT dynamics. Taken together, it seems that SETDB1 can affect various aspects of tumor progression such as cell division and cell migration, at multiple levels by different mechanisms. At the chromatin level, it can affect transcriptional control of key factors to support cell migration and proliferation (Gerlitz, 2020; Orouji et al., 2019; Yu et al., 2019) and it can promote global chromatin condensation to support cell migration (Gerlitz, 2020; Maizels et al., 2017); while at the MT level, it can modulate MT dynamics probably by affecting the activity and/or the binding of MAPs to MTs. Indeed, we were able to identify interaction between SETDB1 and the tubulin deacetylase HDAC6, as well as increased tubulin acetylation levels following SETDB1 KD.

Cross-talk between interphase chromatin and MTs was found before: MT-driven mechanical forces were shown to alter chromatin organization (Gerlitz et al., 2013; Maizels and Gerlitz, 2015), the motor protein kinesin KIF4 was found to be involved in heterochromatin formation, transcription and DNA repair (Mazumdar et al., 2011). Dynein light chain 1 (DLC1) was found to be involved in transcriptional control (Maizels and Gerlitz, 2015). Tau nuclear subcellular localization is thought to associate with DNA damage protection (Diez and Wegmann, 2020). LIS1, a regulator of cytoplasmic dynein was shown to interact with histone H1 and MeCP1 and to affect chromatin binding of the latter (Keidar et al., 2019). Thus, SETDB1 seems to join this growing group of factors with activities in both chromatin and MT worlds.

## Materials and Methods

### Cell culture

B16-F1 cells (ATCC, CRL-6323), WM266.4 cells (a kind gift from Gal Markel, Sheba Medical Center, Israel), HeLa cells (ATCC, CCL-2) and HEK293 cells (ATCC, CRL-1573) were cultured in DMEM (Biological Industries, Kibbutz Beit-Haemek, Israel) supplemented with 10 % fetal calf serum, 0.5 % penicillin -streptomycin solution mix, and 1 % L-glutamine, at 37°C in a 7% CO_2_ environment. Transfections of DNA plasmids were carried out with the NanoJuice Transfection Kit (71900-3, Merck, Kenilworth, NJ, USA) or the Avalanche-Everyday Transfection Reagent (EZT-EVDY-1, EZ Biosystems, College Park, MD, USA) according to the manufacturers’ instructions. Cells were incubated for 24 hours before further analysis. For gene silencing, cells were transfected twice with siRNA (IDT, Coralville, IA, USA), with a time interval of 48 hours using INTERFERin transfection reagent (Polyplus-transfection, Illkirch-Graffenstaden, France) according to the manufacturer’s instructions. Cells were incubated for 24 hours after the second transfection before further analysis. SiRNAs used were mouse SETDB1(mm.Ri.Setdb1.13.1), human SETDB1 (hs.Ri.SETDB1.13.1) and negative control (51-01-14-04). Proliferation rate was measured with Cell Proliferation Kit (XTT based) (20-300-1000, Biological Industries, Kibbutz Beit-Haemek, Israel), according to the manufacturer’s protocol.

### Plasmids and molecular cloning

Plasmids expressing GFP-fused WT and H1224K SETDB1 fused to GFP were generated by PCR using pREV-SETDB1 as a template (a kind gift from Slimane Ait-Si-Ali, Centre National de la Recherche Scientifique (CNRS), Université de Paris, Université Paris Diderot, Paris, France) (Fritsch et al., 2010). WT SETDB1 was amplified by KOD Hot Start DNA Polymerase (71086, Merck) and the oligonucleotides 5’-cagagctcATGTCTTCCCTTCCTG GGTGCAT-3’ and 5’-gtgtcgaCTAAAGAAGACGTCCTCTGCATTCAAT-3’. The PCR product was ligated into SacI-SalI sites in pEGFP-C3. Site directed mutagenesis to generate SETDB1 H1224K was performed by PCR using the above two oligonucleotides and the oligonucleotides: 5’-GGGCCGCTACCTCAACaagAGTTGCAGCCCCAACC-3’ and 5’-GGTTGGGGCTGC AACTcttGTTGAGGTAGCGGCCC-3’. Cloning procedures were confirmed by DNA sequencing. pcDNA-HDAC6-FLAG was a gift from Tso-Pang Yao (Addgene plasmid # 30482) (Kawaguchi et al., 2003). EB1-GFP was a kind gift from Orly Reiner, Weizmann Institute of Science, Rehovot, Israel. pEGFP-Moesin was a kind gift from Peter Vilmos, Biological Research Center of the Hungarian Academy of Sciences, Szeged, Hungary.

### Protein lysate preparation and Western blot analysis

For nuclear–cytoplasmic fractionation, cells were washed in PBS, re-suspended in STM buffer: 50 mM Tris pH 7.4, 250 mM sucrose, 5 mM MgSO_4_, 10 mM NaF, 2 mM DTT, 0.025 % Triton X-100 and 1x protease inhibitor cocktail (539134, Millipore, Burlington, MA, USA) and lysed by a Dounce homogenizer. Following cell membrane breakage (as monitored under the microscope), the nuclei fraction was precipitated at 700 *g* for 10 minutes at 4 °C. The cytoplasmic supernatant was collected to a new tube and the nuclear pellet was re-suspended in TP buffer: 10 mM Tris pH 7.4, 10 mM phosphate buffer pH 7.4, 5 mM MgSO_4_, 10 mM NaF and 1x protease inhibitor cocktail supplemented with 5 % glycerol. Protein concentrations were measured using the Bradford assay. Samples were stored at -80 °C.

For whole cell extracts, cells were washed in PBS, re-suspended in 2x SDS sample buffer (100 mM Tris pH 6.8, 10 % glycerol, 2 % SDS, 100 mM bromophenol blue, 0.1 M DTT and 1x protease inhibitor cocktail) and sonicated. Samples were then heated at 95 °C for 10 minutes and kept at -20 °C until use.

Protein extracts were separated in SDS-PAGE and analyzed by Western blot analysis using the following antibodies: rabbit anti-SETDB1 (1:500, Santa Cruz Biotechnology sc-66884), rabbit anti-SUV39H2 (1:1,000, Abcam ab190270), rabbit anti-histone H3 (1:10,000, Millipore 05-928), mouse anti-α-tubulin (1:5,000, Thermo Fisher Scientific 62204), mouse anti-acetylated tubulin (1:1,000, Santa Cruz Biotechnology sc-23950), mouse anti-β-actin (1:5,000, Sigma-Aldrich A1978), mouse anti-GFP (1:500, Santa Cruz Biotechnology sc-9996) and rabbit anti-FLAG (1:1,000, Sigma-Aldrich F7425). Quantitative data analysis was performed with ImageJ/Fiji software (National Institute of Health, Bethesda, USA).

### Co-immunoprecipitation

HEK293 cells were harvested 48 hours following co-transfection and lysed in extraction buffer (50 mM Tris pH 8, 150 mM NaCl, 20 mM EDTA, 50 mM NaF, 1% Triton X-100) supplemented with 1x protease inhibitor cocktail. Cells were lysed on ice for 30 minutes and centrifuged at 20,000 *g* for 30 minutes at 4°C. A 5% input control sample was taken from each cleared lysate and boiled in SDS sample buffer for 10 minutes at 98 °C. For immunoprecipitation, clarified lysates were supplemented with GFP Trap Magnetic Agarose (gtma-20, Chromotek, Planegg-Martinsried, Germany) and incubated with rotation for 1 hour at 4°C. The beads were washed once in extraction buffer and three times in PBS and boiled in SDS sample buffer for 10 minutes at 98 °C, before being loaded on an 8% acrylamide gel for subsequent Western blot analysis.

### Immunostaining

Cells plated on fibronectin-coated coverslips (03-090-1-05, Biological Industries, Beit-Haemek, Israel) were fixed in methanol supplemented with 1 mM EGTA at -20°C for 6 minutes. Antibodies included rabbit anti-SETDB1 (1:50, Santa Cruz Biotechnology sc-66884), rabbit anti-SETDB1(1:500, Cell signaling Technology 93212), rabbit anti-SUV39H2 (1:120, Abcam ab190270), mouse anti-α-tubulin (1:200, Thermo Fisher Scientific 62204), goat anti-GFP (1:2,000, Abcam ab5450) and rabbit anti-γ-tubulin (1:400, Abcam ab1132). All images were collected using an Olympus 1×81 fluorescent microscope with a coolSNAP HQ2 CCD camera (Photometrics, Tuscon AZ, USA).

### MT recovery assay

Cells plated on fibronectin-coated coverslips were treated with 7 μg·ml^-1^ of nocodazole for 3 hours. Following three washings with cold DMEM to remove the nocodazole, the cells were incubated at 37 °C in pre-warmed complete growth medium for the indicated periods of time, fixed and immunostained as described above. Quantitative data analysis was performed with ImageJ/Fiji software (NHI) by manual delineation of the total area covered by MTOC-linked MTs.

### Time-lapse imaging

For live imaging, cells were plated in 35-mm glass-bottom dishes. Time-lapse images were collected with a coolSNAP HQ2 CCD camera (Photometrics, Tuscon AZ, USA) mounted on an Olympus 1×81 fluorescent microscope at 37°C and 7 % CO_2_. Frames were captured every 3 seconds for 5 minutes (movies to track growing MT ends) and every 3 minutes for 10 hours (movies to monitor mitotic progression). Acquired images were analyzed by ImageJ/Fiji software.

To analyze growing MT ends, MTrackJ plugin was used to track EB1-GFP comets. Comets were analyzed in each frame considering the distal site of the comet as the comet point. Data analysis was done according to duration and length tracked. To analyze mitosis progression, mitotic events were followed in terms of time and success rate.

## Acknowledgments

We thank Slimane Ait-Si-Ali (Centre National de la Recherche Scientifique (CNRS), Université de Paris, Université Paris Diderot, Paris, France), Orly Reiner (Weizmann Institute of Science, Rehovot, Israel) and Peter Vilmos (Biological Research Center of the Hungarian Academy of Sciences, Szeged, Hungary) for providing plasmids, Gal Markel (Sheba Medical Center, Israel) for providing cells, Michal Likvornik and Betsalel Elgrably for calibrating the acetylated tubulin antibody. This research was funded by Ariel University.

## Authors Contributions

Conceptualization, G.G. and R.H.V.; Methodology, R.H.V. and G.G.; Investigation, R.H.V. and N.P.; Formal Analysis, R.H.V.; Writing – Original Draft, G.G.; Writing – Review & Editing, R.H.V., N.P., T.L. and G.G.; Supervision; T.L. and G.G.; Funding Acquisition, G.G.

## Supplementary Figures

**Sup Figure 1.**
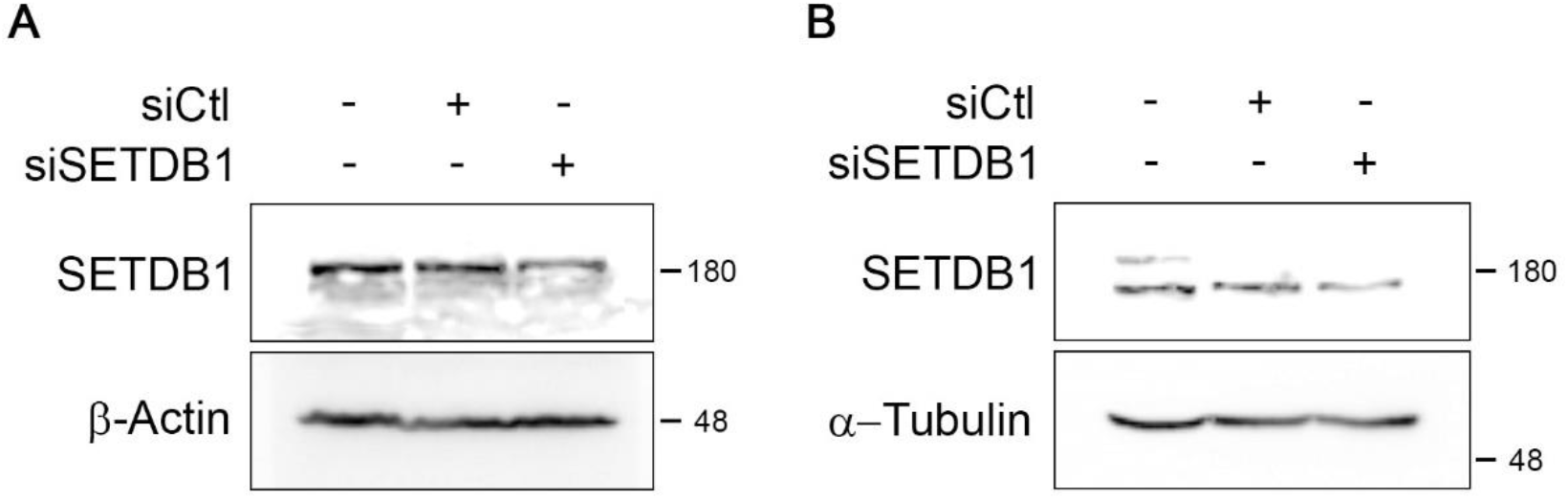
SETDB1 knockdown. **A**,**B** SETDB1 protein levels in mouse B16-F1 cells (A) and in human WM266.4 (B) after treatment with siRNA against SETDB1.

## References

A, M., Latario, C.J., Pickrell, L.E., and Higgs, H.N. (2020). Lysine acetylation of cytoskeletal proteins: Emergence of an actin code. Journal of Cell Biology 219, e202006151.

Batham, Lim, and Rao (2019). SETDB-1: A Potential Epigenetic Regulator in Breast Cancer Metastasis. Cancers 11, 1143.

Beyer, S., Pontis, J., Schirwis, E., Battisti, V., Rudolf, A., Le Grand, F., and Ait-Si-Ali, S. (2016). Canonical Wnt signalling regulates nuclear export of Setdb1 during skeletal muscle terminal differentiation. Cell Discovery 2, 16037.

Bilodeau, S., Kagey, M.H., Frampton, G.M., Rahl, P.B., and Young, R.A. (2009). SetDB1 contributes to repression of genes encoding developmental regulators and maintenance of ES cell state. Genes & Development 23, 2484–2489.

Binda, O., LeRoy, G., Bua, D.J., Garcia, B.A., Gozani, O., and Richard, S. (2010). Trimethylation of histone H3 lysine 4 impairs methylation of histone H3 lysine 9: Regulation of lysine methyltransferases by physical interaction with their substrates. Epigenetics 5, 767–775.

Ceol, C.J., Houvras, Y., Jane-Valbuena, J., Bilodeau, S., Orlando, D.A., Battisti, V., Fritsch, L., Lin, W.M., Hollmann, T.J., Ferre, F., et al. (2011). The histone methyltransferase SETDB1 is recurrently amplified in melanoma and accelerates its onset. Nature 471, 513–517.

Chen, K., Zhang, F., Ding, J., Liang, Y., Zhan, Z., Zhan, Y., Chen, L., and Ding, Y. (2017). Histone Methyltransferase SETDB1 Promotes the Progression of Colorectal Cancer by Inhibiting the Expression of TP53. J. Cancer 8, 3318–3330.

Chin, H.G., Esteve, P.-O., Ruse, C., Lee, J., Schaus, S.E., Pradhan, S., and Hansen, U. (2020). The microtubule-associated histone methyltransferase SET8, facilitated by transcription factor LSF, methylates α-tubulin. Journal of Biological Chemistry 295, 4748–4759.

Cho, S., Park, J.S., and Kang, Y.-K. (2013). Regulated nuclear entry of over-expressed Setdb1. Genes Cells 18, 694–703.

Cruz-Tapias, P., Zakharova, V., Perez-Fernandez, O., Mantilla, W., Ramírez-Clavijo, S., and Ait-Si-Ali, S. (2019). Expression of the Major and Pro-Oncogenic H3K9 Lysine Methyltransferase SETDB1 in Non-Small Cell Lung Cancer. Cancers 11, 1134.

Cuellar, T.L., Herzner, A.-M., Zhang, X., Goyal, Y., Watanabe, C., Friedman, B.A., Janakiraman, V., Durinck, S., Stinson, J., Arnott, D., et al. (2017). ---Silencing of retrotransposons by SETDB1 inhibits the interferon response in acute myeloid leukemia--. The Journal of Cell Biology 216, 3535–3549.

Diez, L., and Wegmann, S. (2020). Nuclear Transport Deficits in Tau-Related Neurodegenerative Diseases. Front. Neurol. 11.

Dodge, J.E., Kang, Y.-K., Beppu, H., Lei, H., and Li, E. (2004). Histone H3-K9 Methyltransferase ESET Is Essential for Early Development. Molecular and Cellular Biology 24, 2478–2486.

Du, D., Katsuno, Y., Meyer, D., Budi, E.H., Chen, S., Koeppen, H., Wang, H., Akhurst, R.J., and Derynck, R. (2018). Smad3-mediated recruitment of the methyltransferase SETDB1/ESET controls Snail1 expression and epithelial–mesenchymal transition. EMBO Rep 19, 135–155.

Fei, Q., Shang, K., Zhang, J., Chuai, S., Kong, D., Zhou, T., Fu, S., Liang, Y., Li, C., Chen, Z., et al. (2015). Histone methyltransferase SETDB1 regulates liver cancer cell growth through methylation of p53. Nat Commun 6, 8651.

Fritsch, L., Robin, P., Mathieu, J.R.R., Souidi, M., Hinaux, H., Rougeulle, C., Harel-Bellan, A., Ameyar-Zazoua, M., and Ait-Si-Ali, S. (2010). A Subset of the Histone H3 Lysine 9 Methyltransferases Suv39h1, G9a, GLP, and SETDB1 Participate in a Multimeric Complex. Molecular Cell 37, 46–56.

Gauchier, M., Kan, S., Barral, A., Sauzet, S., Agirre, E., Bonnell, E., Saksouk, N., Barth, T.K., Ide, S., Urbach, S., et al. (2019). SETDB1-dependent heterochromatin stimulates alternative lengthening of telomeres. Sci. Adv. 5, eaav3673.

Gerlitz, G. (2020). The Emerging Roles of Heterochromatin in Cell Migration. Front. Cell Dev. Biol. 8, 394.

Gerlitz, G., Reiner, O., and Bustin, M. (2013). Microtubule dynamics alter the interphase nucleus. Cell. Mol. Life Sci. 70, 1255–1268.

Guo, J., Dai, X., Laurent, B., Zheng, N., Gan, W., Zhang, J., Guo, A., Yuan, M., Liu, P., Asara, J.M., et al. (2019). AKT methylation by SETDB1 promotes AKT kinase activity and oncogenic functions. Nat Cell Biol 21, 226–237.

Hachiya, R., Shiihashi, T., Shirakawa, I., Iwasaki, Y., Matsumura, Y., Oishi, Y., Nakayama, Y., Miyamoto, Y., Manabe, I., Ochi, K., et al. (2016). The H3K9 methyltransferase Setdb1 regulates TLR4-mediated inflammatory responses in macrophages. Scientific Reports 6, 28845.

Ho, Y., Lin, Y., Huang, Y., Chang, J., Yeh, K., Lin, L., Gong, Z., Tzeng, T., and Lu, J. (2017). Significance of histone methyltransferase SETDB 1 expression in colon adenocarcinoma. APMIS 125, 985–995.

Hwang, Y.J., Han, D., Kim, K.Y., Min, S.-J., Kowall, N.W., Yang, L., Lee, J., Kim, Y., and Ryu, H. (2014). ESET methylates UBF at K232/254 and regulates nucleolar heterochromatin plasticity and rDNA transcription. Nucleic Acids Res. 42, 1628–1643.

Jiang, Y., Jakovcevski, M., Bharadwaj, R., Connor, C., Schroeder, F.A., Lin, C.L., Straubhaar, J., Martin, G., and Akbarian, S. (2010). Setdb1 Histone Methyltransferase Regulates Mood-Related Behaviors and Expression of the NMDA Receptor Subunit NR2B. Journal of Neuroscience 30, 7152–7167.

Jiang, Y., Loh, Y.-H.E., Rajarajan, P., Hirayama, T., Liao, W., Kassim, B.S., Javidfar, B., Hartley, B.J., Kleofas, L., Park, R.B., et al. (2017). The methyltransferase SETDB1 regulates a large neuron-specific topological chromatin domain. Nat Genet 49, 1239–1250.

Jurkowska, R.Z., Qin, S., Kungulovski, G., Tempel, W., Liu, Y., Bashtrykov, P., Stiefelmaier, J., Jurkowski, T.P., Kudithipudi, S., Weirich, S., et al. (2017). H3K14ac is linked to methylation of H3K9 by the triple Tudor domain of SETDB1. Nat Commun 8, 2057.

Kang, Y.-K. (2018). Surveillance of Retroelement Expression and Nucleic-Acid Immunity by Histone Methyltransferase SETDB1. BioEssays 40, 1800058.

Kang, Y.-K., and Min, B. (2020). SETDB1 Overexpression Sets an Intertumoral Transcriptomic Divergence in Non-small Cell Lung Carcinoma. Front. Genet. 11, 573515.

Karanth, A.V., Maniswami, R.R., Prashanth, S., Govindaraj, H., Padmavathy, R., Jegatheesan, S.K., Mullangi, R., and Rajagopal, S. (2017). Emerging role of SETDB1 as a therapeutic target. Expert Opinion on Therapeutic Targets 21, 319–331.

Karimi, M.M., Goyal, P., Maksakova, I.A., Bilenky, M., Leung, D., Tang, J.X., Shinkai, Y., Mager, D.L., Jones, S., Hirst, M., et al. (2011). DNA Methylation and SETDB1/H3K9me3 Regulate Predominantly Distinct Sets of Genes, Retroelements, and Chimeric Transcripts in mESCs. Cell Stem Cell 8, 676–687.

Kawaguchi, Y., Kovacs, J.J., McLaurin, A., Vance, J.M., Ito, A., and Yao, T.P. (2003). The deacetylase HDAC6 regulates aggresome formation and cell viability in response to misfolded protein stress. Cell 115, 727– 738.

Keidar, L., Gerlitz, G., Kshirsagar, A., Tsoory, M., Olender, T., Wang, X., Yang, Y., Chen, Y.-S., Yang, Y.-G., Voineagu, I., et al. (2019). Interplay of LIS1 and MeCP2: Interactions and Implications With the Neurodevelopmental Disorders Lissencephaly and Rett Syndrome. Front. Cell. Neurosci. 13, 370.

Keniry, A., Gearing, L.J., Jansz, N., Liu, J., Holik, A.Z., Hickey, P.F., Kinkel, S.A., Moore, D.L., Breslin, K., Chen, K., et al. (2016). Setdb1-mediated H3K9 methylation is enriched on the inactive X and plays a role in its epigenetic silencing. Epigenetics & Chromatin 9, 16.

Kostaki, M., Manona, A.D., Stavraka, I., Korkolopoulou, P., Levidou, G., Trigka, E.-A., Christofidou, E., Champsas, G., Stratigos, A.J., Katsambas, A., et al. (2014). High-frequency p16 ^INK 4A^ promoter methylation is associated with histone methyltransferase SETDB1 expression in sporadic cutaneous melanoma. Experimental Dermatology 23, 332–338.

Lawson, K.A., Teteak, C.J., Zou, J., Hacquebord, J., Ghatan, A., Zielinska-Kwiatkowska, A., Fernandes, R.J., Chansky, H.A., and Yang, L. (2013). Mesenchyme-specific Knockout of ESET Histone Methyltransferase Causes Ectopic Hypertrophy and Terminal Differentiation of Articular Chondrocytes. Journal of Biological Chemistry 288, 32119–32125.

Li, L., and Yang, X.-J. (2015). Tubulin acetylation: responsible enzymes, biological functions and human diseases. Cell. Mol. Life Sci. 72, 4237–4255.

Liu, S., Brind’Amour, J., Karimi, M.M., Shirane, K., Bogutz, A., Lefebvre, L., Sasaki, H., Shinkai, Y., and Lorincz, M.C. (2014). Setdb1 is required for germline development and silencing of H3K9me3-marked endogenous retroviruses in primordial germ cells. Genes Dev. 28, 2041–2055.

Loyola, A., Bonaldi, T., Roche, D., Imhof, A., and Almouzni, G. (2006). PTMs on H3 Variants before Chromatin Assembly Potentiate Their Final Epigenetic State. Molecular Cell 24, 309–316.

Maizels, Y., and Gerlitz, G. (2015). Shaping of interphase chromosomes by the microtubule network. FEBS Journal 282, 3500–3524.

Maizels, Y., Elbaz, A., Hernandez-Vicens, R., Sandrusy, O., Rosenberg, A., and Gerlitz, G. (2017). Increased chromatin plasticity supports enhanced metastatic potential of mouse melanoma cells. Exp. Cell Res. 357, 282–290.

Matsui, T., Leung, D., Miyashita, H., Maksakova, I.A., Miyachi, H., Kimura, H., Tachibana, M., Lorincz, M.C., and Shinkai, Y. (2010). Proviral silencing in embryonic stem cells requires the histone methyltransferase ESET. Nature 464, 927–931.

Mazumdar, M., Sung, M.-H., and Misteli, T. (2011). Chromatin maintenance by a molecular motor protein. Nucleus 2, 591–600.

Minkovsky, A., Sahakyan, A., Rankin-Gee, E., Bonora, G., Patel, S., and Plath, K. (2014). The Mbd1-Atf7ip-Setdb1 pathway contributes to the maintenance of X chromosome inactivation. Epigenetics Chromatin 7, 12.

Mozzetta, C., Boyarchuk, E., Pontis, J., and Ait-Si-Ali, S. (2015). Sound of silence: the properties and functions of repressive Lys methyltransferases. Nature Reviews Molecular Cell Biology 16, 499–513.

Orouji, E., Federico, A., Larribère, L., Novak, D., Lipka, D.B., Assenov, Y., Sachindra, S., Hüser, L., Granados, K., Gebhardt, C., et al. (2019). Histone methyltransferase SETDB1 contributes to melanoma tumorigenesis and serves as a new potential therapeutic target. Int. J. Cancer 145, 3462–3477.

Park, I.Y., Powell, R.T., Tripathi, D.N., Dere, R., Ho, T.H., Blasius, T.L., Chiang, Y.-C., Davis, I.J., Fahey, C.C., Hacker, K.E., et al. (2016). Dual Chromatin and Cytoskeletal Remodeling by SETD2. Cell 166, 950–962.

Rodriguez-Paredes, M., Martinez de Paz, A., Simó-Riudalbas, L., Sayols, S., Moutinho, C., Moran, S., Villanueva, A., Vázquez-Cedeira, M., Lazo, P.A., Carneiro, F., et al. (2014). Gene amplification of the histone methyltransferase SETDB1 contributes to human lung tumorigenesis. Oncogene 33, 2807–2813.

Schultz, D.C. (2002). SETDB1: a novel KAP-1-associated histone H3, lysine 9-specific methyltransferase that contributes to HP1-mediated silencing of euchromatic genes by KRAB zinc-finger proteins. Genes & Development 16, 919–932.

Sharif, J., Endo, T.A., Nakayama, M., Karimi, M.M., Shimada, M., Katsuyama, K., Goyal, P., Brind’Amour, J., Sun, M.-A., Sun, Z., et al. (2016). Activation of Endogenous Retroviruses in Dnmt1 −/− ESCs Involves Disruption of SETDB1-Mediated Repression by NP95 Binding to Hemimethylated DNA. Cell Stem Cell 19, 81–94.

Strepkos, D., Markouli, M., Klonou, A., Papavassiliou, A.G., and Piperi, C. (2020). Histone methyltransferase SETDB1: a common denominator of tumorigenesis with therapeutic potential. Cancer Res canres.2906.2020.

Sun, Q.-Y., Ding, L.-W., Xiao, J.-F., Chien, W., Lim, S.-L., Hattori, N., Goodglick, L., Chia, D., Mah, V., Alavi, M., et al. (2015). SETDB1 accelerates tumourigenesis by regulating the WNT signalling pathway. J. Pathol. 235, 559–570.

Tachibana, K., Gotoh, E., Kawamata, N., Ishimoto, K., Uchihara, Y., Iwanari, H., Sugiyama, A., Kawamura, T., Mochizuki, Y., Tanaka, T., et al. (2015). Analysis of the subcellular localization of the human histone methyltransferase SETDB1. Biochemical and Biophysical Research Communications 465, 725–731.

Takikita, S., Muro, R., Takai, T., Otsubo, T., Kawamura, Y.I., Dohi, T., Oda, H., Kitajima, M., Oshima, K., Hattori, M., et al. (2016). A Histone Methyltransferase ESET Is Critical for T Cell Development. J. Immunol. 197, 2269–2279.

Torrano, J., Al Emran, A., Hammerlindl, H., and Schaider, H. (2019). Emerging roles of H3K9me3, SETDB1 and SETDB2 in therapy-induced cellular reprogramming. Clin Epigenetics 11, 43.

Tsusaka, T., Shimura, C., and Shinkai, Y. (2019). ATF7IP regulates SETDB1 nuclear localization and increases its ubiquitination. EMBO Rep. e48297.

Wang, G., Long, J., Gao, Y., Zhang, W., Han, F., Xu, C., Sun, L., Yang, S.-C., Lan, J., Hou, Z., et al. (2019). SETDB1-mediated methylation of Akt promotes its K63-linked ubiquitination and activation leading to tumorigenesis. Nat Cell Biol 21, 214–225.

Wong, C.-M., Wei, L., Law, C.-T., Ho, D.W.-H., Tsang, F.H.-C., Au, S.L.-K., Sze, K.M.-F., Lee, J.M.-F., Wong, C.C.-L., and Ng, I.O.-L. (2016). Up-regulation of histone methyltransferase SETDB1 by multiple mechanisms in hepatocellular carcinoma promotes cancer metastasis. Hepatology 63, 474–487.

Yang, L., Mei, Q., Zielinska-Kwiatkowska, A., Matsui, Y., Blackburn, M.L., Benedetti, D., Krumm, A.A., Taborsky, G.J., and Chansky, H.A. (2003). An ERG (ets-related gene)-associated histone methyltransferase interacts with histone deacetylases 1/2 and transcription co-repressors mSin3A/B. Biochemical Journal 369, 651–657.

Yu, L., Ye, F., Li, Y.-Y., Zhan, Y.-Z., Liu, Y., Yan, H.-M., Fang, Y., Xie, Y.-W., Zhang, F.-J., Chen, L.-H., et al. (2019). Histone methyltransferase SETDB1 promotes colorectal cancer proliferation through the STAT1-CCND1/CDK6 axis. Carcinogenesis bgz131.

Zeng, T.-B., Han, L., Pierce, N., Pfeifer, G.P., and Szabó, P.E. (2019). EHMT2 and SETDB1 protect the maternal pronucleus from 5mC oxidation. Proc Natl Acad Sci USA 116, 10834–10841.

Zhang, Y., Huang, J., Li, Q., Chen, K., Liang, Y., Zhan, Z., Ye, F., Ni, W., Chen, L., and Ding, Y. (2018). Histone methyltransferase SETDB1 promotes cells proliferation and migration by interacting withTiam1 in hepatocellular carcinoma. BMC Cancer 18, 539.

